# Optical induction of autophagy via *Transcription factor EB* (TFEB) reduces pathological tau in neurons

**DOI:** 10.1101/600908

**Authors:** JL Binder, V Deretic, JP Weick, K Bhaskar

**Affiliations:** Department of Molecular Genetics and Microbiology, University of New Mexico, Albuquerque, NM 87131, USA; Department of Neurosciences, University of New Mexico, Albuquerque, NM 87131, USA

## Abstract

Aggregation and accumulation of microtubule associated protein tau in neurons is major neuropathological hallmark of Alzheimer’s disease (AD) and related tauopathies. Attempts have been made to promote clearance of pathological tau (p-Tau) from neurons via autophagy. Over expression of transcription factor EB (TFEB), has shown to clear pTau from neurons via autophagy. However, sustained TFEB activation and/or autophagy can create burden on cellular bioenergetics and can be deleterious. Thus, we engineered a minimally invasive optical system that could transiently alter autophagic flux. We optimized and tested an optogenetic gene expression system derived from a previously engineered bacterial transcription factor, EL222. For the first time, our group utilized this system not only to spatial-temporally control nuclear TFEB expression, we also show light-induced TFEB has the capacity to reduce p-Tau burden in AD patient-derived human iPSC-neurons. Together, these results suggest that optically-regulatable gene expression of TFEB unlocks opto-therapeutics to treat AD and other dementias.

## Introduction

Among the various microtubule-associated proteins (MAP), tau (encoded by *MAPT*) predominately localizes to axons where it binds to microtubules. Tau is known to promote nucleation, stabilization, and prevent disassembly of microtubules^1^. However, tau is susceptible to many post-translational modifications^2^, with phosphorylation being one of the well-studied modifications^3,4,5^. Upon hyperphosphorylation, tau linearly decreases its affinity to microtubules causing depolymerization^6^, which has been the prevailing hypothesis, that tau’s loss-of-function causes neurodegeneration ^7,8^. These dissociated forms of tau can self-assemble into paired-helical filaments (PHFs) gaining further potential to aggregate as Neurofibrillary tangles (NFTs) – a classical neuropathological hallmark of Alzheimer’s disease (AD) and related tauopathies^9^. Alternatively, hyperphosphorylated and pathological tau (p-Tau) has been shown to acquire gain-of-toxic function in triggering synaptotoxicity relevant to AD^10^. While AD is the most common form of tauopathy and sixth leading cause of death in the United States^11^, NFT pathology is also the primary etiology in many, but rare tauopathies such as Progressive Supranuclear Palsy (PSP), Pick’s disease (PiD), Corticobasal Degeneration (CBD), Fronto-temporal Dementia and Parkinsonism linked to Chromosome-17 tau-type (FTDP-17T) and others^12^. Because of the exponential rise in tauopathy related deaths, there is an urgent need to find intervention(s) against tauopathies.

A plausible strategy to prevent p-Tau from becoming pathological is to promote its degradation via autophagy in “*at risk*” neuronal populations. Impairment of autophagic processes has been implicated in several neurodegenerative disorders. However, there are still several reasons for considering autophagy’s potential in clearing p-Tau: (i) Multiple reports have suggested a causal link between failure in autophagic processing and AD-related pathologies^13,14,15,16,17,18,19^, (ii) Autophagy disposes of potentially toxic intracellular protein aggregates too large for proteasomal removal^20,21^, and performs trophic functions^22,23^ (sometimes referred to as “programmed cell survival”), (iii) autophagy removes depolarizes mitochondria^24^ and shows cytoprotective interactions with stressed endoplasmic reticulum^25,26,27,28,29,30^ that are of direct consequence for tauopathies^31,32,33,34,35,36^, (iv) previous work from our group provides compelling evidence that autophagy prevents spurious inflammasome activation^37,38^, which when left uncontrolled, could drive tau pathology and cognitive impairment^39^, and (v) chronic promotion of autophagic processing can enhance clearance of p-Tau and rescue neurotoxicity in a mouse model of tauopathy^40^. We have demonstrated that induction of autophagy via chemical or genetic means lead to clearance of inflammation-induced p-Tau^41^ in neuronal cells. Transcription Factor EB (TFEB) regulates transcription of an entire CLEAR (Coordinated Lysosomal Expression and Regulation) network, which consists of a consensus site predominately found in the promoter regions of autophagy-lysosomal genes^42,43^. Thus, when TFEB localization is nuclear, it leads to robust increase in lysosome biogenesis, and results in accelerated degradation of autophagic substrates ^30,44,45,46^. Phosphorylation of Ser211 in TFEB by mammalian target of rapamycin complex 1 or mechanistic target of rapamycin complex 1 (mTORC1) is one of the key regulators of nuclear localization, as the pS211 phosphorylation prevents TFEB from entering into the nucleus^47^. However, the limitation of pro-autophagy studies is their focus on the continual activation of autophagy. While autophagy is generally thought to promote survival as discussed above, under certain conditions sustained autophagic-flux can lead to cell death^48,48^. Furthermore, prolonged activation of autophagy proteins (e.g. LC3 and BECN1) and vacuoles in response to ischemic stroke/reperfusion *in vivo*, or oxygen-glucose deprivation (OGD) *in vitro* leads to significant cell death^49^. Interestingly many autophagic processes do not significantly affect cell health until days after the injury, indicating that prolonged activation is critical for cell death to occur^50,51^. Another example, constitutive activation of the δ2 glutamate receptor was demonstrated to cause Purkinje cell death in Lurcher mice via activation of autophagy^52^. Thus, for elderly tauopathy patients with co-morbid conditions such as ischemia and vascular dementia, sustained activation of autophagy could exacerbate cell death rather than reduce it. Therefore, it is important to develop tunable systems to turn-on/turn-off autophagy in neurons with optimum spatio-temporal control.

In the case of autophagy, which requires the coordinated expression and function of a host of proteins, optical induction of transcription by a master regulator offers such a system^53^. Here we optimized an optical induction system based on light-responsive bacterial transcription factor^52^ to drive TFEB expression in a number of different mammalian expression systems that display pathological tau. For the first time, our group has shown that optically controlled TFEB efficiently expresses in human AD neurons, up-regulates TFEB target genes, and efficiently targets and reduces pTau.

## RESULTS

### Cytomegalovirus (CMV) promoter and the nuclear localization signal (NLS) sequence derived from cMyc shows robust gene expression with light

We chose a light-inducible gene expression system that utilizes an engineered version of EL222, a bacterial transcription factor that contains a Light-Oxygen-Voltage (LOV) protein^72,54^(reiterated in Fig. 1A). This system was previously optimized with five copies of the bacterial EL222-binding Clone 1–20 base pairs (C120)_5_ sequence^55,56^. This consensus site acts like a promoter region for the EL222 binding and drives the expression of any genes inserted downstream of C120 (FIG 1A-B).

**Figure 1.**
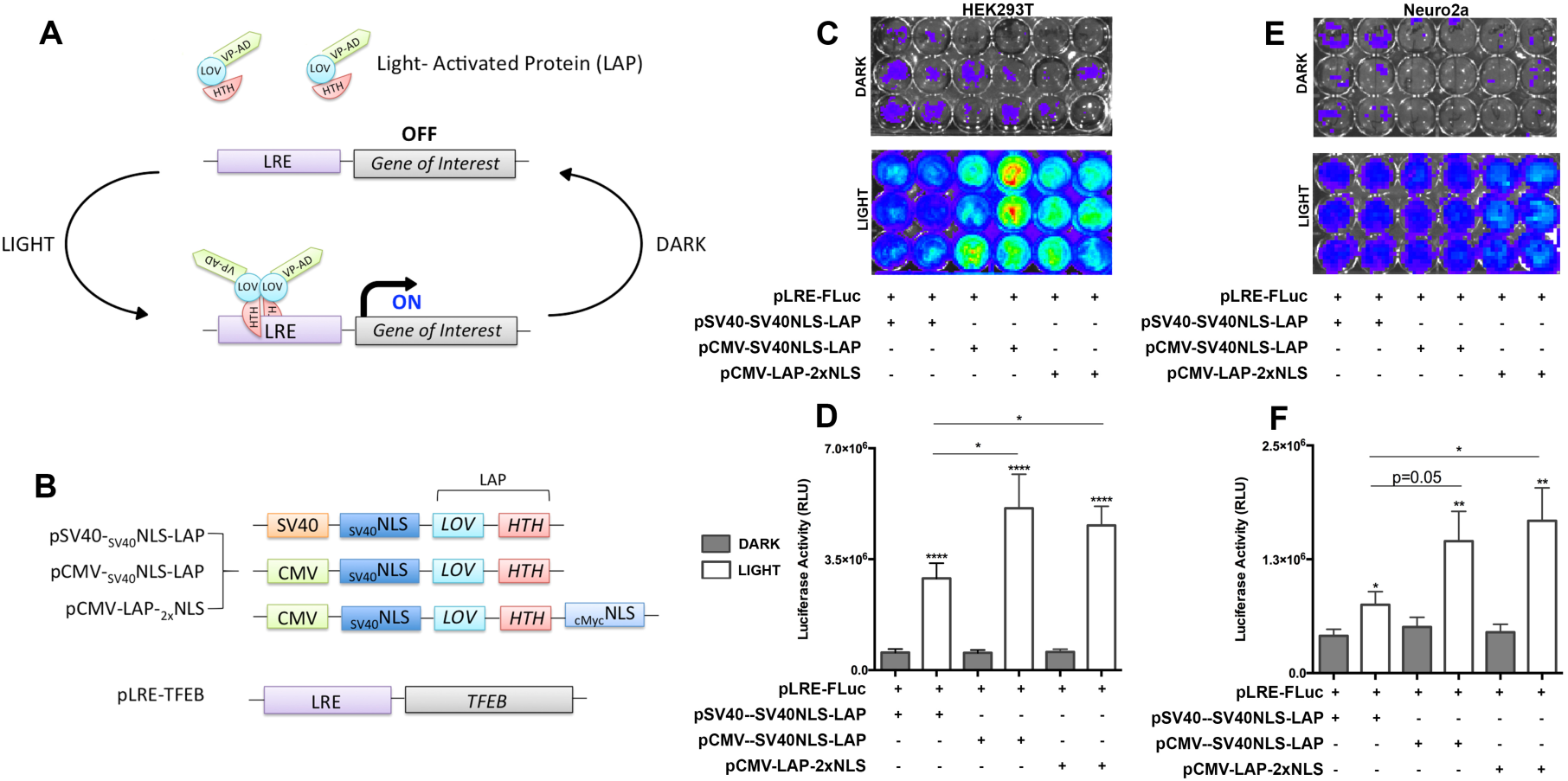
Optogenetic gene expression system in neuronal cell line. **A.** Schematic of previously established gene expression system derived from an EL222 bacterial transcription factor, termed Light-Activated Protein (LAP). **B**. Schematic of our optimized changes made to the LAP construct for successful neuronal transfection/induction as well as TFEB cloned into the LRE construct. **C-D**. Quantitative comparison of various versions of LAP constructs using pLRE-Firefire Luciferase reporter, (pLRE-FLuc) in <u>HEK293T’s</u> measuring luciferase activity units (RLU) via radiance levels detected by IVIS (mean ±s.e.m, Student’s *t* test or one-way ANOVA with Tukey multiple comparison test, ****p<0.0005 n=5) **E-F.** Quantitative comparison of various versions of LAP constructs using pLRE-FLuc in Neuroblastoma cell line <u>N2a’s</u> measuring Luciferase activity units via radiance levels detected by IVIS. (mean ±s.e.m, Student’s *t* test or one-way ANOVA with Tukey multiple comparison test, **p<0.05 n=5)

In order to recapitulate Motta-Mena et al., EL222 system, we first tested their original two-plasmid system pVP-EL222, (that we call ‘light activated protein’ or ‘LAP’); and their pC120-Fluc (Firefly Luciferase reporter that we call ‘light-response element’ or ‘LRE’), where the luciferase gene was inserted downstream of LRE. As reported previously^72^, we also observed robust luciferase activation upon blue light illumination in HEK293T cells (Fig. 1C-D). However, using the original pSV40-_SV40_NLS-LAP in our neuro2a (N2a) cell line, we show more than a two-fold decrease in luciferase expression compared to HEK293Ts. Therefore we created two different EL222 light-responsive systems by replacing the SV40 promoter with a CMV promoter and including an additional cMyc nuclear localization signal (NLS) sequence (Fig. 1B). Strikingly, we observe statistically significant, over two-to four-fold luciferase expression upon blue-light stimulation compared to dark controls in both HEK293T and N2a’s (Fig. 1C-F). We also observed a different degree of luciferase expression with different promoter and NLS combinations with the pCMV-LAP-_2x_NLS showing the robust induction of luciferase expression in N2a cells. This difference was likely due to a weaker SV40 promoter in the LAP plasmid used in the original system^72^ compared to more robust CMV promoter, as previously reported^57^. Our observations in N2a cells also aligned with the robustness of CMV promoter in previously reported in N2a’s cells^58^. Furthermore, we also reasoned that the NLS from SV40 might have contributed to the poor induction efficiency, as the addition of cMyc NLS to the N-Terminus^59^ in ‘pCMV-LAP-_2x_NLS’ showed the highest induction of luciferase activity in both HEK293T and N2a cells (FIG 1B-F). Because of the robust induction efficiency of pCMV-LAP-_2x_NLS, we used only this plasmid in the subsequent experiments.

### Constitutively active TFEB (not light inducible) clears various types of pathological tau with equal efficiency in cellular models of tauopathy

Now that the optogenetic system parameters have been set, there are a few other factors that need to be considered within our neuronal tauopathy model. First, as mentioned above, there are many forms of tauopathy. We wanted to determine whether TFEB has the ability to target different forms of pathological tau (pTau) and lead to its clearance via autophagy in neuronal cells. Tau gene (*MAPT*) in humans encodes six different isoforms that are contrasted by exons 2, 3 and 10 ^60^. Exon 10 encodes second microtubule binding repeat, thereby resulting in tau with either three (3R – without second repeat)-or four (4R – with second repeat) microtubule binding repeats of 31–32 amino acids in the carboxy terminal half ^85^. Tau isoforms are also generated in either having or not having one (1N), two (2N), or zero (0N) amino terminal inserts of 29 amino acids each in the N-terminal half of the protein^85^. In normal adult brain, the relative amounts of 3R tau and 4R tau are approximately equal, however in neurodegenerative tauopathies, the ratio of 3R:4R is often altered ^61^. Besides altered isoform rations, post-translational modifications, such as phosphorylation in tau, can also affect tau’s function and contribute to disease pathogenesis in tauopathies^86^. We and others have previously shown that by TFEB degrades pTau via beclin-1 dependent autophagy pathway^31,32^. However, the effectiveness of TFEB on different isoforms of tau with different disease-modifications has not been tested. To test this, we co-transfected TFEB with different forms of tau; **(1)** pCMV-0N3R – non-mutant tau, but over-expression can lead to Pick’s Disease (PiD) ^62^; **(2)** pCMV-0N3R(T231D/S235D), which mimics hyperphosphorylation on T231/S335 sites and known to disrupt tau’s interaction with tubulins^63^ **(3)** pCMV-0N4R P301L, which causes FTDP-17T^64,65^; **(4)** pCMV-0N4R – non-mutant tau, but over-expression can lead to progressive supranuclear palsy (PSP)^66^. Co-transfection of TFEB with different types of tau lead to consistent reduction in all types of overexpressed total tau levels in N2a cells (Fig. 2A-B), with T231D/S235D phosphorylation-mimicking tau showing the most significant reduction (Fig. 2A-B). Together, these results suggest that TFEB can clear different types of pTau with robust consistency in neuronal cells. Furthermore, since T231 residue can acquire potent neurotoxic conformation called *cis*-pTau (or ‘Cistauosis’, as a result of phosphorylation of tau at T231)^67,68,69^,TFEB’s role in significantly reducing T231D/S235D species of pTau suggest the potential therapeutic potential of targeting TFEB against tauopathies.

**Figure 2.**
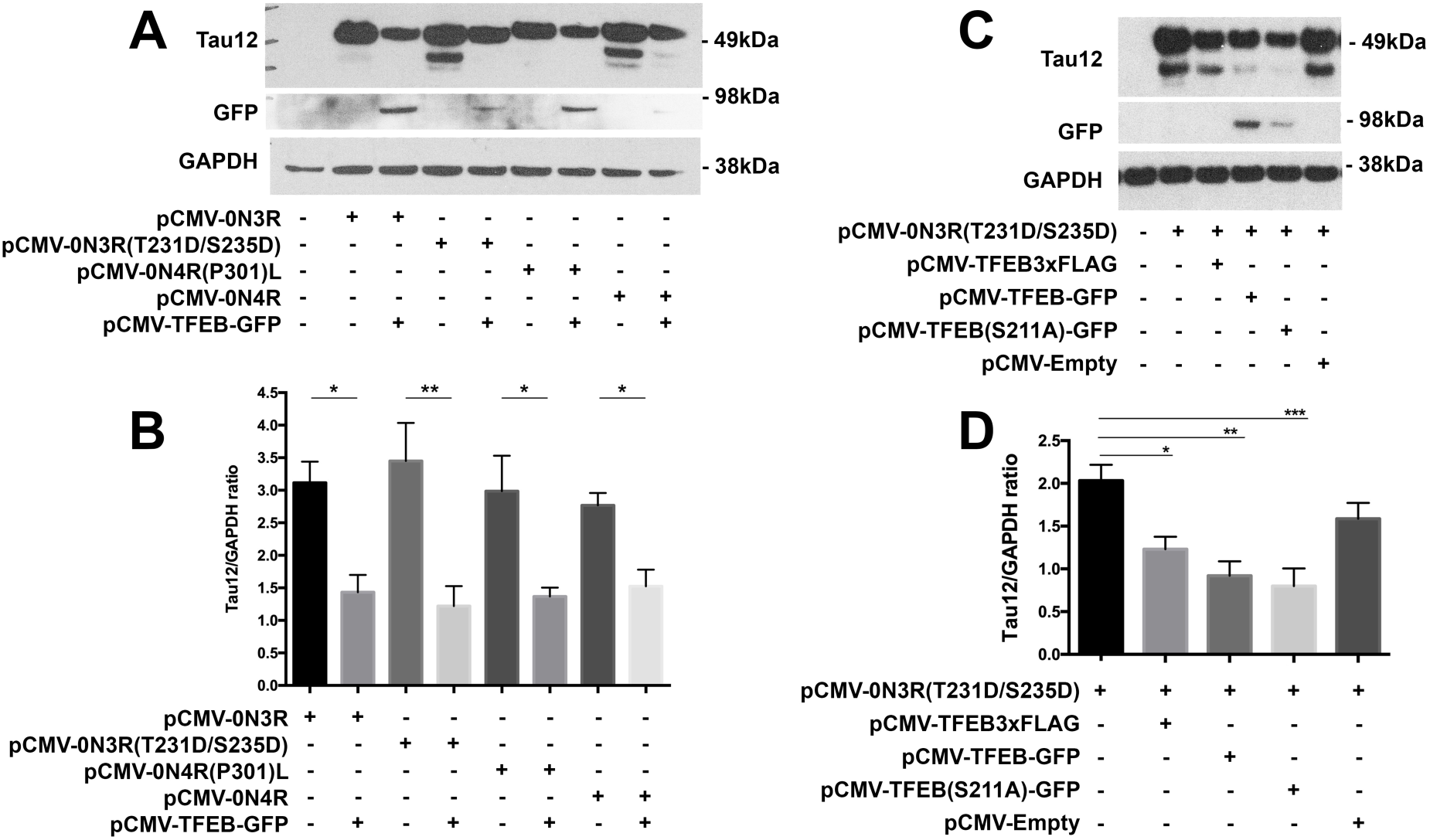
TFEB differentially targets various forms of pTau. **A-B.** Western blot and quantification showing significant reduction in various forms of tau via WT – 0N3R, (0N3R) T231D/S235D, (0N4R) P301L, and WT – 0N4R with the addition of constitutive overexpression of TFEB activity. Results indicated most forms of tau are equivalently reduced by TFEB, however (0N3R) T231D/S235D shows highest significance in expression and reduction. Total tau/GAPDH ratio (mean ±s.e.m, Student’s *t* test, *\*\**p<0.01 n=3) **C-D.** Western blot and quantification showing significantly reduced (0N3R) T231D/S235D with the addition of various forms of constitutive TFEB overexpression; pCMV-TFEB3xFLAG, pCMV-TFEB-GFP, pCMV-TFEB(S211A)GFP. Results indicate pCMV-TFEB(S211A)GFP holds the best yield in total tau reduciton. Total tau/GAPDH ratio (mean ±s.e.m, one-way ANOVA with Tukey multiple comparison test, ***p<0.005 n=4) **E-F.** With the addition of Bafilomycin, western blot and quantification showing an increased trend in LC3-II levels in pCMV-TFEB(S211A)GFP. LC3-II/GAPDH ratio (mean ±s.e.m, one-way ANOVA with Tukey multiple comparison test, n=3)

Next, we determine whether or not different tags (FLAG versus GFP) to the TFEB have any differential effects in clearing pTau via autophagy. Co-expression of T231D/S235D tau with either pCMV-TFEB3xFLAG or pCMV-TFEB-GFP showed that GFP tagged TFEB was slightly efficient in clearing mutant tau than 3xFLAG tagged TFEB (Fig. 2C-D). Given that TFEB has to be nuclear for it to be functionally efficient and to drive the expression of genes in the CLEAR network, we next assessed the effects of S211 phosphorylation in TFEB in clearing mutant tau. TFEB-GFP with the S211A mutation, which prevents phosphorylation by mTORC1^70^ thus promotes TFEB’s nuclear entry, had the greatest effect in reducing T231D/S235D mutant pTau (FIG2 C-D). Together, these results suggest that genetically facilitating the nuclear entry of TFEB does provide added advantage in enhancing the autophagic clearance of T231D/S235D tau.

### Optogenetically expressed TFEB activates CLEAR network genes in neuronal cells

To determine the optimal parameters for the optogenetically-driven TFEB (Opto-TFEB) expression in N2a cells, we co-transfected N2a cells with the two different LAP’s and pLRE-TFEB(S211A)GFP plasmids, stimulated the cells with blue light for 12 h, fixed in 4% PFA, immunostained for VP16 and GFP and performed confocal microscopy. Double immunofluorescence analysis (for VP16 and GFP), with the substitution of a CMV promoter and additional cMyc NLS, we observed a significant increase of TFEB expression (as revealed by anti-GFP staining) with blue-light stimulation compared to dark control (Fig. 3A-B). As expected, the VP16 staining was detectable in pCMV-LAP (Fig. 3A).

**Figure 3.**
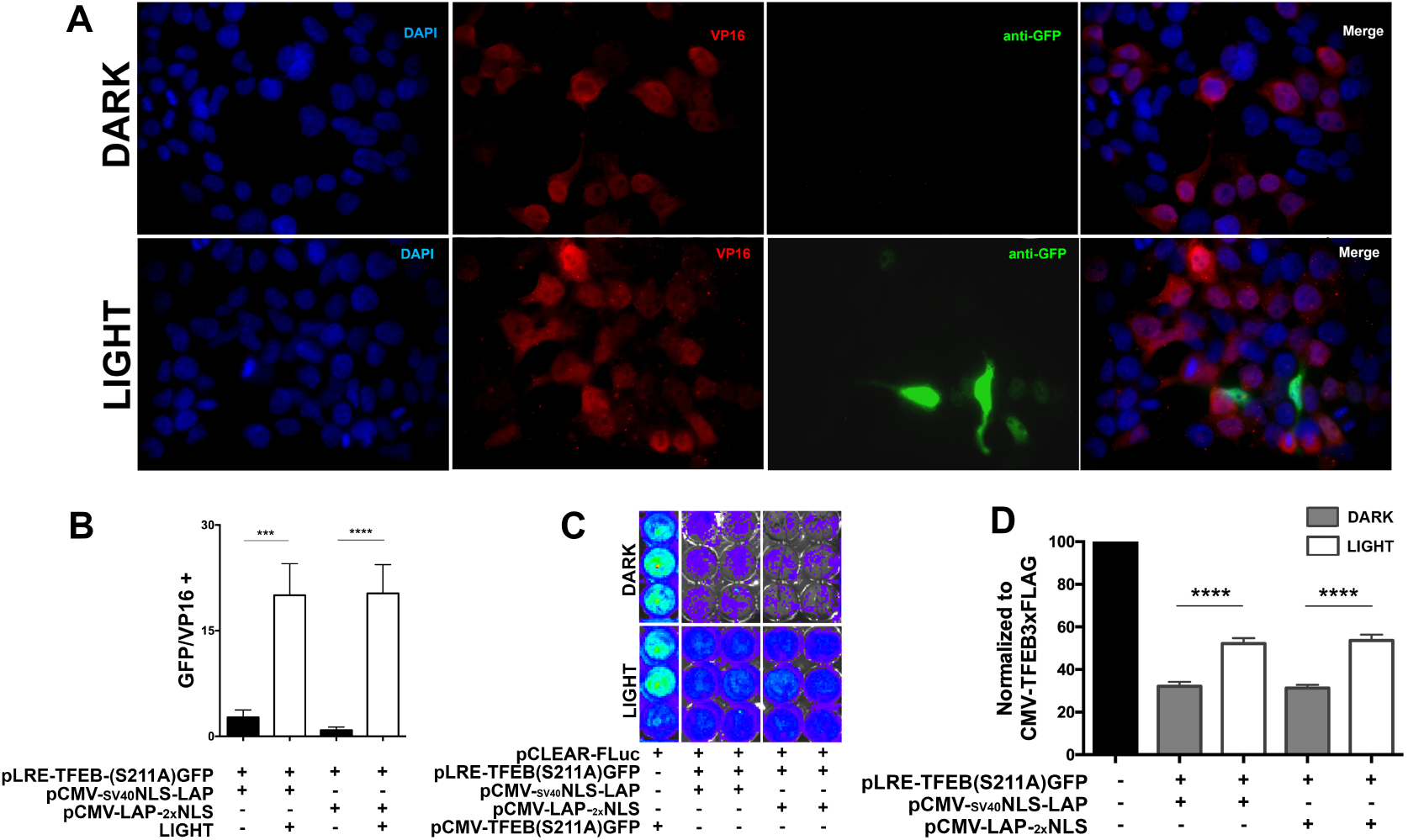
Optogenetic TFEB induction in neuronal cell line and CLEAR activity readout. **A-B.** Quantitative immunocytochemistry showing significant increase in TFEB expression in Light control vs Dark, comparison of various versions of LAP constructs using pLRE-TFEB-(S211A)GFP (mean ±s.e.m, Student’s *t* test, ****p<0.0005, n=4). **C-D**. Quantitative comparison of various versions of LAP constructs using pCLEAR-Firefly Luciferase reporter, (pCLEAR-Fluc) in <u>N2a’s</u> measuring luciferase activity units (RLU) via radiance levels detected by IVIS (mean ±s.e.m, Student’s *t* test or one-way ANOVA with Tukey multiple comparison test, ****p<0.0005 n=4)

After having established that TFEB expression can be induced optogenetically (Opto-TFEB) in neuronal cells, we set to determine whether light-induced TFEB expression is functionally active and display expected transcriptional function. Based on the TFEB’s preferential binding to the CLEAR consensus site (^5’^GTCACGTGAC^3’^), we next utilized a previously published^71^, reporter plasmid pCLEAR-FLuc to assess the functional activation of Opto-TFEB. The pCLEAR-FLuc plasmid consists of four replicates of the CLEAR consensus sequence upstream of the luciferase gene, serving as a DNA-binding activity readout for TFEB. We transiently co-transfected pCLEAR-FLuc with pLAPs, and pLRE-TFEB(S211A)GFP in N2a cells and stimulated the cells with blue light overnight (12 h). Shortly after luciferin treatment, whole cell culture plates were imaged using IVIS Lumina Series III. We observed a significant increase in TFEB DNA-binding activity (enhanced CLEAR-luciferase signal) only in cells that optogenetically expressed TFEB with light, and not in the dark control (Fig.3C-D). Together, our results suggest that Opto-TFEB is functionally active in driving expected transcriptional activity of genes in the CLEAR network.

### Opto-TFEB reduces pathological tau in neuronal cells

Because we have shown Opto-TFEB can drive transcriptional regulation of the CLEAR network (responsible for autophagy flux and lysosomal biogenesis), then it should in theory increase the autophagic flux and reduce the levels of misfolded p-Tau upon light stimulation. We took a multiple approach in establishing that Opto-TFEB is capable of clearing specifically phosphorylated species of Tau. First, we used confocal microscopy to measure the levels of pTau in N2a cells that expressed Opto-TFEB. We overexpressed human mutated 0N3R-T231D/S2345D tau along with pCMV-LAP2xNLS and pLRE-TFEB(S211A)GFP. Upon cell-by-cell analysis, we observed that for every Opto-TFEB^±^cell there is little to no Tau12 staining (Fig. 4A). Using Zeiss software to measure total geometrical intensity levels we saw significant increase in Opto-TFEB levels compared to dark. To perform an unbiased quantification, we performed quantitative morphometry for Tau12 levels using high-content and automated Cellomics® microscopy. We observed a significant decrease in overall Tau12 intensity levels in Opto-TFEB^+^ cells compared to dark control (FIG4 C-D). Next, to validate the Cellomics® data, we performed Western blot analysis for total human tau (Tau12) in N2a cells. Strikingly, we observed statistically significant increase in TFEB-GFP expression (Fig. 4D-E) and reduction in the levels of total tau (Tau12) (Fig. 4D-F) in samples with Opto-TFEB (+Light) compared to controls Opto-TFEB (Dark). Together, these results suggest that light-induced expression of TFEB is capable of reducing pathological (T231D/S235D) tau in N2a cell line.

**Figure 4.**
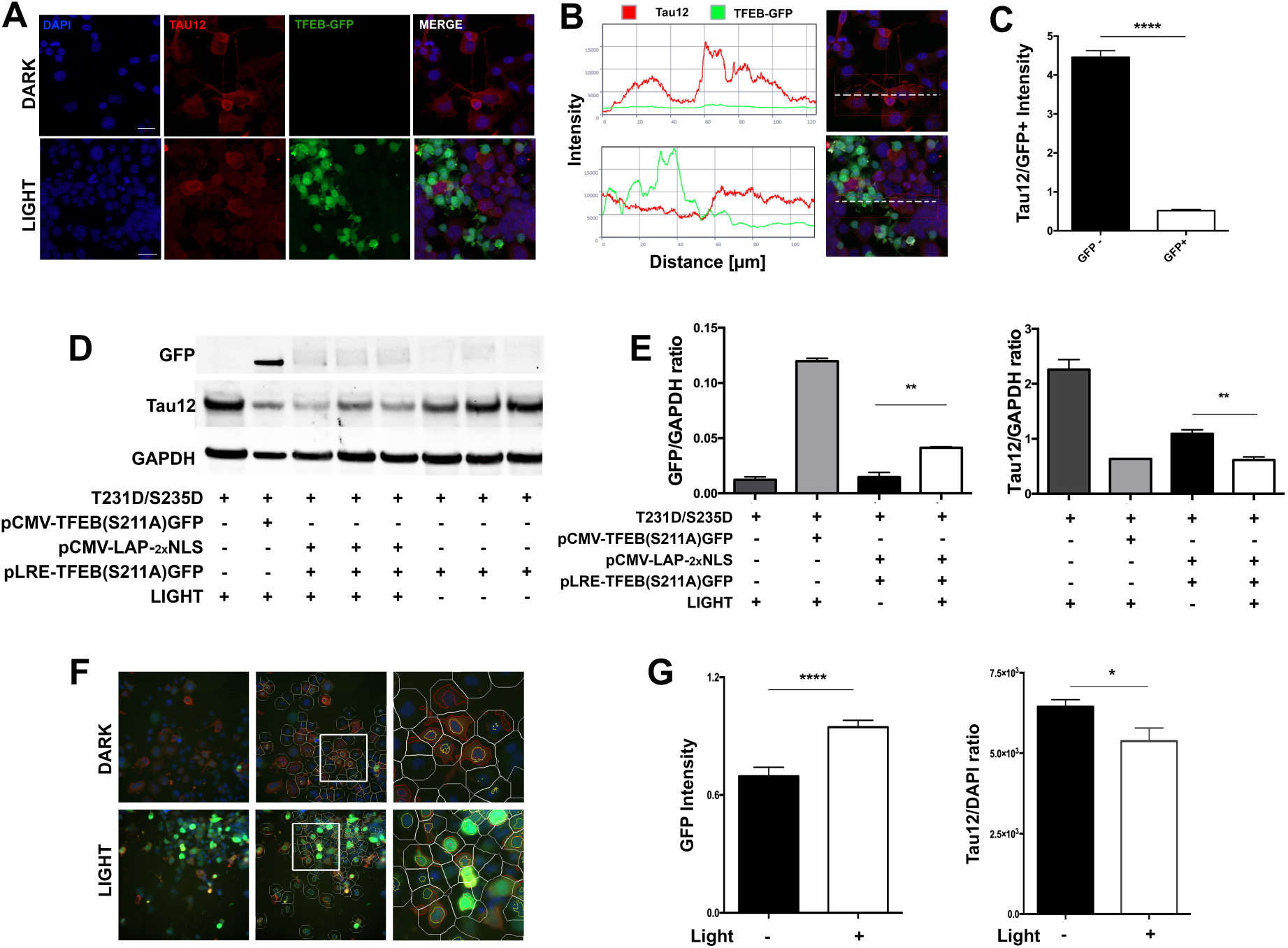
Optogenetic TFEB induction in neuronal cell line reduces neuronal pathological mimicking tau. **A.** Quantitative confocal immunocytochemistry using N2a cells overexpressing human 0N3R-T231D/S235D tau show lack of colocalization of optogenetically induced TFEB expression with Tau12 positive cells. **B.** Representation of colocalization profile for Tau12 (red) and LRE-TFEB(S211A)GFP (green) analysis **C.** Quantification was done using histo-profile analysis via ZEISS ZEN imaging Software. (mean ±s.e.m, Student’s t test, *p<0.05, n=3). **D-E**. Western Blot analysis showing overall total protein levels are reduced when Opto-TFEB is expressed via light stimulation compared to dark. **F-G.** Cellomics®-based high-content imaging analysis of the effects on total Tau levels within Dark and Light controls. Cells were automatically identified based on nuclear staining (DAPI), then cells were selected for positive nuclear green fluorescence (TFEB(S211A)GFP) to further analyze for Tau12 (RED) intensity levels within 100 pixel radius per cell. Briefly, white lines represent cell boundaries, red lines represent positive cytosolic Tau12, and yellow lines indicate nuclear TFEB(S211A)GFP-positive cells, then subjected by automated image analysis. Scale bars: 10 µm. Quantitative morphometric data (mean ±s.e.m, Student’s t test, *p<0.05, n=3)

**Figure 5.**
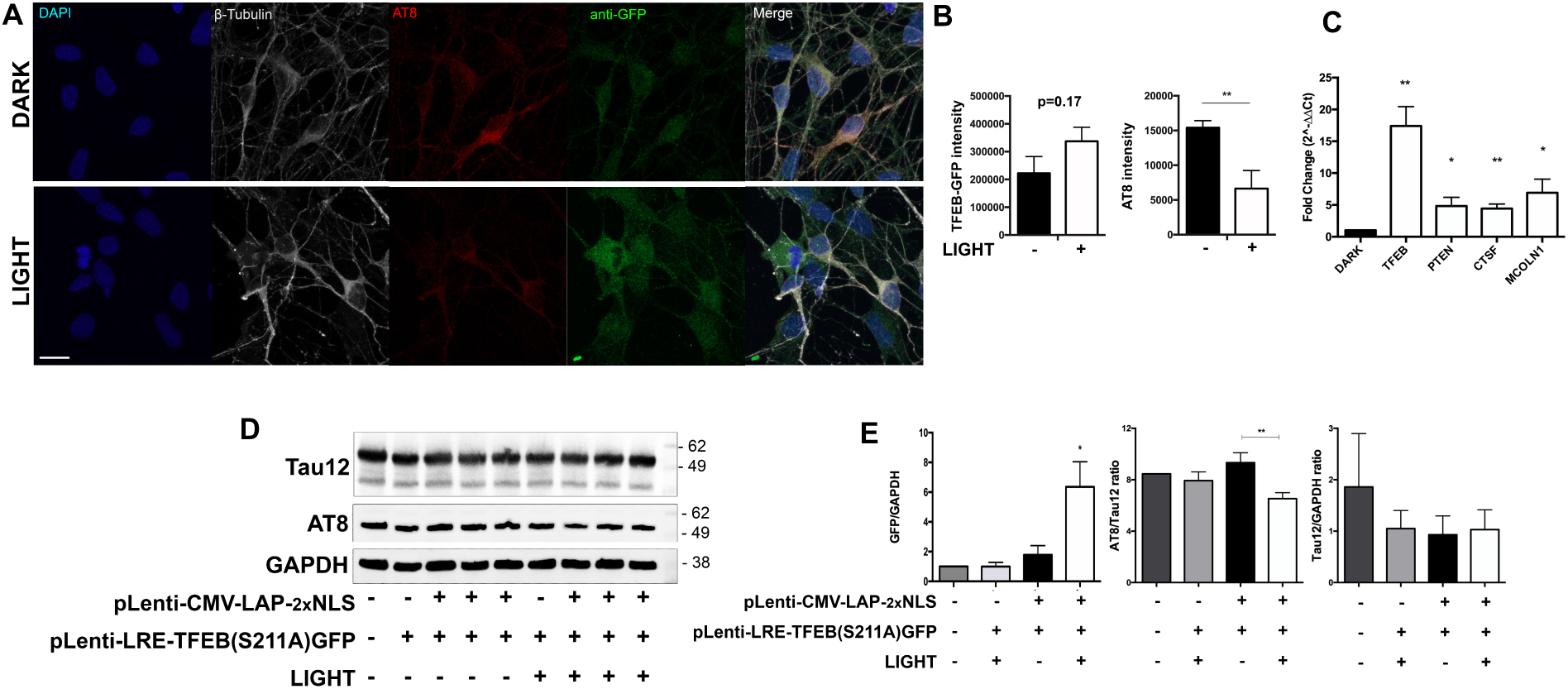
Optogenetic TFEB clears pTau in human induced pluripotent stem cells derived into neurons (iPSNs). **A-B.** Quantitative immunocytochemistry showing significant increase in TFEB expression with subsequent lower levels of pTau (AT8) in Light control compared to Dark, using viral-particle versions, pGF1-CMV-LAP-2xNLS and pGF1-LRE-TFEB-(S211A)GFP (mean ±s.e.m, Student’s t test, ****p<0.0005, n=3). **C.** RT-qPCR analysis of TFEB gene expression and TFEB targets (PTEN, CTSF, and MCOLN1). Each sample was taken 24 hours of subsequent 12 hour light stimulation. **D-E.** Western blot and quantification showing significantly reduced pMAPT (AT8 and AT180) with the transduction of viral optogenetic TFEB and subsequent light stimulation. (mean ±s.e.m, Student’s t test, *p<0.05, n=3)

While our results, using N2a cell model for the first time, demonstrates that Opto-TFEB expression can reduce p-Tau, this is an artificially overexpression system in neuroblastoma cell line and may not necessarily represent a true disease model of neurodegenerative tauopathies. Moreover, we objectively overloaded the system by competing three different transgene (pRC-CMV-0N3R(T231D/S2345D) tau, pCMV-LAP_2x_NLS, and pLRE-TFEB(S211A)GFP) expression plasmids. It has been shown that multi-transfection efficiency diminishes in a hyperbolic manner^72^; consequently the percentage of tri-positive transfected cell is lowered. Unfortunately, this makes it difficult to analyze true representation of reduction efficacy for our transient, but regulatable optogenetic approach.

Therefore, to compensate for the possible energetically competing transgene expression, we next tested the efficacy of Opto-TFEB in a previously characterized AD relevant iPSCs cell line called ‘sAD2.1’^73^. In our hands, sAD2.1 iPSC neurons (iPSNs), display robust pTau (positive for AT8, AT180, and PHF1) levels (supplementary FIG).

To assess the efficacy of Opto-TFEB in sAD2.1 cells, it was necessary to clone Opto-TFEB plasmids (pCMV-LAP2xNLS and pLRE-TFEB(S211A)GFP) into a lentiviral system and subsequently co-transduce sAD2.1 iPSNs. Using confocal microscopy and analyzing with Zeiss software for overall geometric intensities, we note there is a significant decrease in pTau (AT8) levels in our light control compared to dark (FIG5 A-B). We also wanted to make sure that the reduction is due strictly because of Opto-TFEB induction. To do so, we measured the gene expression levels of three known TFEB targets; PTEN^38^, CTSF, and MCOLN1^40^. (FIG5 C) Compared to Dark, we show significant increase in TFEB expression as well as the three targets we chose. Lastly, we took a look at overall levels of pTau (AT8 and AT180) at the total protein level (FIG5 D-E). In conclusion, for the first time we show that light controlled / optogenetic TFEB has the ability to reduce pTau in a highly respectable tauopathy model.

## DISCUSSION

TFEB is a master transcriptional regulator of autophagy and lysosome biogenesis. However, studies have revealed TFEB constitutes and interacts with a variety of biological functions, including the inflammatory process, stress responsive pathways^74^, oxidative stress^75^, and metabolic regulation^76^. It has been shown in many studies how autophagy contributes to human AD diseased iPSNs and the endogenous phenotypes/disease characteristics. For example, Reddy et al. generated iPSC-derived human forebrain cortical neurons from AD patients with M146L and A246E mutations, as well as a presenilin-1 (PS-1) knockdown in control neurons^77^. Using the same CLEAR-luciferase reporter assay, they found a reduction in CLEAR activity in their forebrain cortical iPSNs as well as decrease in LC3II levels in PS-1 knockdown neurons, advocating decreased autophagy flux. Another group, again used iPSC-derived forebrain cortical neurons studied the amino acid metabolite homocysteine (Hcy), and found that exposure to Hcy caused reduced autophagic activity by a means of increase in mTORC1 activity therefore reduction in TFEB activity by phosphorylation of TFEB by mTORC1 ^78^.

Therefore, in order to setup a system that employs TFEB as a therapeutic target, it cannot be constitutively active or termed in an active/nuclear state. Here we describe a new blue light inducible TFEB gene expression system that works well in mouse neuronal cell lines and human AD iPSCs derived into mature neurons. Not only have we shown successful light controlled gene expression, but we also effectively enhanced the autophagy flux, specifically targeting and reducing pathological tau in our AD iPSNs. The benefits of using diseased iPSNs in this study, is that when derived into a neuronal population, the AD phenotype is displayed ^98,79,80,81^. This avoids the hurdle of low tri-plasmid transfection efficiency.

Now that we have successfully shown the benefits of neuronal optoTFEB induction, it may be interesting to see what remunerations could be achieved when cell-type specific optoTFEB is accomplished. This would answer the question if it poses more beneficial characteristics if optoTFEB only was induced in glia cells, like astrocytes or microglia, using a GFAP or CX3CR1 promoter to drive the LAP transgene. It also may be necessary to induce OptoTFEB in various time-points, to assess cell toxicity and or the ability of OptoTFEB to be beneficiary in a dose-like dependent manner. Altogether, our data strongly suggest that OptoTFEB efficiently expresses in AD iPSNs, up-regulates TFEB target genes, and efficiently targets and clears pTau.

## METHODS

### Vector construction

All constructs were cloned using NEB HIFI Assembly Kit (NEB # E5520S) with restriction enzymes and PCR amplification. Briefly, the original episomal plasmids gifted by Motta-Mena, (pVP-EL222 and pGL4-C120-mCherry) were cloned into different backbones with subsequent promoters and /or gene of interest; pN1-CMV-TFEB-GFP (Addgene # 38119). Newly cloned episomal plasmids were then additional cloned into lentivector backbone, pGF1-NfkB-EF1-Puro (Systemsbio # TR012PA-P). Q5® Site-Directed Mutagenesis Kit was used to make mutations (S142A and S211A) in TFEB gene (NEB # E0445S). All Tau constructs used; 1) pRC/CMV -0N3R-tau (human tau with three microtubule-binding repeats with no N-terminal inserts); 2) 0N4R-tau (human tau with four microtubule-binding repeats with no N-terminal inserts); 3) 0N4R-P301L (human tau with four microtubule-binding repeats with P301L FTDP-17T mutation); 4) 0N3R-T231D/S235D. See supplemental Table 1 for all cloned vectors and their corresponding names.

### Cell Lines

#### HEK293T and Neuro-2a

(ATCC # CRL-3216 and #CCL-131, respectively) cells were maintained at 37°C in 5% CO2 in DMEM supplemented with 10% FBS, 5% penicillin/streptomycin, and grown in 24-well plates. For transient transfections, cells were split the day before ∼ 1-4 × 10^5^ cells/well, therefore 70-80% confluence the following day. Before transfection, media was replaced with phenol red free media, (FluoroBrite DMEM; ThermoFisher # A1896701). Cells were then transfected with Lipofectamine 2000 (Invitrogen) as per company’s protocol. Dilutions of various plasmid concentrations were as followed for a 24-well plate; pLAP’s – [2000ng/µL], pLRE’s – [500ng/µL], pCMV-TFEB’s – [500ng/µL], pCMV-hTau’s – [1000ng/µL], pCLEAR-FLuc – [500ng/µL]. Therefore always, a 1:4 ratio of LRE to LAP.

### Induced pluripotent stem cells

**sAD2.1; Coriell # GM24666,** *(iPSCs from Fibroblast NIGMS Human Genetic Cell Repository Description: ALZHEIMER DISEASE; AD Affected: Yes. Gender: Male. Age: 83 YR (At Sampling). Race: Caucasian.)*

Briefly, iPSCs were maintained in mTESR +supplement (StemCell # 85850) Neuron differentiation followed the StemCells neuronal differentiation kit/protocol; (StemCell #05835, #05833, #08500, #08510). Later medium was changed to BrainPhys™ without Phenol Red (StemCell #05791) for optical induction. (Neural progenitor cells seeded at 1.5x 10^4^ cells/cm^2^ for maturation)

### Light Induction

12hrs post transfection, an in-house blue LED device (465 nm, strip of LEDs glued to PCB board; Amazon) was placed 8cm or 16cm above the plate. Note, the constraints of the light source also had to be altered (twice the distance than our cell lines; 16cm) due to higher sensitivity of iPSNs to the blue-light and the heat it produces, compared to N2a cell lines. The intensity of the light received by cells was measured to be to 8 W/m^2^; as per suggestion of Motta-Mena et al.) Verified, using the LI-190 Quantum Sensor and LI-250A light meter (LI-COR Biosciences). The LED strips were connected to SLBSTORES 3528 5050 12V DC Mini Remote Controller (Amazon) for variations of on/off patterns to best match a cycle of 20 s on and 60 s off as recommended per Motta-Mena et al. The control plate was kept in a PCB blackout box with breathable air slots, (a shelf in the incubator, above and away from the light source shelf). For transiently transfected cells, 24hrs post-transfection, samples were collected/fixed for analysis.

### Lentivirus production and luciferase assay

Using HEK293T’s, seeded in 100mm plates. Lentiviral Transgenes were cloned into the pGF1-EF1-Puro backbone. Lentiviral packaging vectors: pMD.2, pPAX2 (Invitrogen cat. no. K4975-00). Cells were transfected with plasmid mix using CaPO_4_ precipitation method, as per protocol Tiscornia et al. 2006 Nat Protocols “Production and purification of lentiviral vectors”. After 48-hrs interval, the viral supernatant was then filtered through 0.45 µm membranes and mixed overnight with cat#631232 Lenti-X™ Concentrator. The next day, samples were centrifuged at 1,500 x g for 45 minutes at 4°C. An off-white pellet is then resuspended in subsequent media, ex: if iPSNs, then neurobasal. Lentiviral titer was measured using cat#631280 Lenti-X™ GoStix™ Plus. Lentiviral Transduction on iPSNs – an IFU of 1×10^6/mL were added to the neurons to make ∼MOI = 2. We transduced sAD2.1 neural progenitor cells 24 hours after plating on poly-ornithine/laminin coated coverslips following StemCell maturation protocol. Subsequently, two weeks after transduction, (Day 40) iPSNs are subjected to light stimulation (12 hours) or kept in the dark, samples were then collected/fixed for analysis.

For Firefly luciferase activities, 4XCLEAR-luciferase reporter plasmid #66800, purchased from addgene. D-Luciferin, Potassium Salt (ThermoFisher # L2916) was reconstituted in water and was added (1:100) to each well, 3-4 mins after addition of substrate, 24-well plate samples were analyzed through the IVIS Lumina Series II with system software.

### Western blotting (WB) and immunocytochemistry (ICC)

#### WB

Cells were lysed by RIPA buffer (Thermo #89900), incubated on ice for 30 mins then centrifuged at 20,000 × g for 15 min. Cell lysate supernatants were then sonicated for 20secs at 30%, then subjected to SDS-PAGE usage, transferred to PVDF membranes and detected using the ECL method (Pierce). Protein levels were quantified using ImageJ (National Institute of Health). Antibodies included; tau12, GAPDH, FLAG, GFP, TFEB, AT8, AT180, LC3B, LAMP1.

#### ICC

Cells were plated on coverslips coated with laminin, once cells were ready for fixation, they were fixed in 4% PFA, blocked with 0.2%triton and 10% donkey serum, incubated in primary overnight in 4°C (5% DS), secondaries were incubated for 1hr at RT. Incubated in DAPI for 10 mins, and mounted to slides using fluoromount(CAT#). Immunofluorescence confocal microscopy was carried out using Zeiss LSM 510 Meta microscope. Histo and profile analysis was performed using ZEISS ZEN imaging Software.

### Gene expression analysis

RNA from cells was extracted using the TriZOL reagent as described by the manufacturer (Thermo Fisher Scientific). Total RNA (20 ng/µL) was converted to cDNA using the High Capacity cDNA Reverse Transcription kit (Thermo Fisher Scientific) and amplified using specific TaqMan assays (catalog # 4331182; Thermo Fisher Scientific). GAPDH (catalog # 4352339E, Thermo Fisher Scientific) was used as a housekeeping gene for normalization. qRT-PCR assays were run on the StepOnePlus® Real-Time PCR System (Thermo Fisher Scientific) and the statistical analyses were performed using Prism.

### Cellomics®-based high-content imaging analysis

Cells were plated in 96 well plates transiently transfected with pCMV-T231D/S235D (phosopho-mimicking tau), pCMV-LAP2xNLS, and pLRE-TFEB(S211A)GFP. Twenty-four hours later, cells were incubated with conditioned medium from BV2’s, as previously described, then subsequently induced with light (470nm) for 12 hours. Cells were fixed in 4% PFA, blocked with 0.2%triton and 10% donkey serum, incubated in primary for one hour at RT (5% DS), secondary was incubated for 1hr at RT. Incubated in DAPI for 10 mins and analyzed through cellomics machine.

### Statistics

Unless otherwise indicated, comparisons between the two groups were done via unpaired t test; comparisons between multiple treatment groups were done via one-way or two-way analysis of variance (ANOVA) with indicated multiple comparisons post-hoc tests. All statistical analyses were performed using GraphPad Prism®.

|  |  |  |
| --- | --- | --- |
| **Plasmids used:** |  |  |
| **Motta-Mena et al** | pGL4-SV40-VP-EL222 | Bacteria Transcription Factor, EL222, LOV domain. |
| **Motta-Mena et al** | pC120-MCH | mCherry reporter |
| **Motta-Mena et al** | pC120-FLuc | Firefly Luciferase reporter |
| **Addgene #38119** | pN1-CMV-TFEB-GFP | Constitutive TFEB- GFP reporter |
| **Binder et al.** | pN1-CMV-TFEB(S211A)-GFP | Constitutive TFEB with (S211A) mutation- GFP reporter |
| **Light response element** | pN1-LRE-TFEB3xFLAG WT | LRE-Flag reporter (data not shown) |
|  | pN1-LRE-TFEB(S142A)3xFLAG | LRE-Flag reporter (data not shown) |
|  | pN1-LRE-TFEB-GFP WT | LRE-GFP reporter (data not shown) |
|  | pN1-LRE-TFEB(S211A)-GFP | LRE-GFP reporter |
| **Light activated protein** | pN1-CMV-EL222 | LAP, CMV promoter, Sv40 NLS N term |
|  | pN1-CMV-EL222-<sub>2x</sub>NLS | LAP, CMV promoter, Sv40-NLS, and cMyc NLS |
| **Lentiviral constructs** | pGF1-CMV-EL222-<sub>2x</sub>NLS | Lenti-LAP |
|  | pGF1-LRE-TFEB(S211A)-GFP | Lenti-LRE-TFEB-GFP reporter |

<sup>1</sup> Kadavath H, Hofele RV, Biernat J, Kumar S, Tepper K, Urlaub H, Mandelkow E, Zweckstetter M. Tau stabilizes microtubules by binding at the interface between tubulin heterodimers. *Proc Natl Acad Sci U S A*.;112(24):7501-6. doi:10.1073/pnas.1504081112. (2015).
<sup>2</sup> Ludovic Martin, Xenia Latypova, Faraj Terro. Post-translational modifications of tau protein: Implications for Alzheimer's disease. *Neurochem Int*. 2011 Mar;58(4):458-71. doi: 10.1016/j.neuint.2010.12.023. (2011)
<sup>3</sup> Hernández F, García-García E, Avila J. Microtubule depolymerization and tau phosphorylation. *J Alzheimers Dis* 37(3):507-13. doi: 10.3233/JAD-130545 (2013)
<sup>4</sup> Gong CX1, Iqbal K. Hyperphosphorylation of microtubule-associated protein tau: a promising therapeutic target for Alzheimer disease. *Curr Med Chem*. 2008;15(23):2321-8. (2008)
<sup>5</sup> Wang JZ, Xia YY, Grundke-Iqbal I, Iqbal K. Abnormal hyperphosphorylation of tau: sites, regulation, and molecular mechanism of neurofibrillary degeneration. *J Alzheimers Dis*. 2013;33 Suppl 1:S123-39. (2013)

